# Ex-*Lactobacillus* Strains with Intrinsic Propensity to Stabilize Pickering Oil-in-Water Emulsions

**DOI:** 10.1101/2023.05.13.540633

**Authors:** Musemma Kedir Muhammed, Xiaoyi Jiang, Elhamalsadat Shekarforoush, Kathryn Whitehead, Finn K. Vogensen, Jens Risbo, Nils Arneborg

**Affiliations:** University of Copenhagen, Department of Food Science, Rolighedsvej 30, DK-1958, Copenhagen, Denmark; Manchester Metropolitan University, Department of Life Sciences, Chester St, Manchester, M15GD, United Kingdom

**Keywords:** bacteria, emulsion, hydrophobicity, ex-*Lactobacillus*, Pickering, stabilization, zeta potential

## Abstract

Knowledge of surface characteristics is a major step in the evaluation of bacterial cells for potential use as Pickering emulsion stabilizers. Here, the cell surface characteristics of 31 strains of the ex-*Lactobacillus* genus were studied with the aim of evaluating their intrinsic abilities to serve as Pickering stabilizers of oil-in-water emulsions. About 77.42% of the tested strains demonstrated relatively highly negative zeta potential (-43.76 mV ≤ zeta potential ≤ -19.23 mV), while ∼58% of the strains demonstrated high cell surface hydrophobicity (microbial adhesion to hexadecane or MATH ≥ 30%). By combining these findings, four different cell surface features were defined (I, II, II and IV). Strains mainly demonstrated the type I surface feature (∼45%), with most expressing strongly negative zeta potential and high surface hydrophobicity (zeta potential < -15 mV and MATH ≥ 30%, respectively). It appeared that the abundance of negative charge on the surfaces of ex-*Lactobacillus* cells positively influences surface hydrophobicity. Assessment of intrinsic Pickering stabilization potential using 12 selected strains indicated that four strains showed profound droplet size stability. At least one strain was observed to have natural propensity to form relativley compact and small emulsion droplets (63±3 µm), leading to enhanced firmness and storage stability of the Pickering emulsions.

## Introduction

Oil-in-water emulsions are known to be unstable and Pickering stabilization can often provide better stability than low molecular surfactants. Pickering particles can be defined as partly hydrophobic solid particles, smaller in size than air bubbles or droplets which possess high binding energies, thus providing irreversible adsorption to interfaces and hence enhanced emulsion stability. Fat crystal particles, for instance, act as Pickering particles in a few food products, such as butter and margarine [1]. Pickering emulsion stability depends on emulsion droplet size, which results from complex interplay between, among others, the ratio of the dispersed and continuous phases and particle size and concentration [2]. The mechanisms of physical instability of emulsions include flocculation, coalescence, creaming and Ostwald ripening [3], [4]. The alterations in these physical mechanisms affect the spatial distribution and structural organization of the molecules [3], [5].

There is renewed interest towards using bio-based materials, such as bacterial particles, for the stabilization of emulsions. According to a few studies, hydrophobic bacteria can stabilize oil-in-water emulsions for an extended period, and bacteria with intermediate hydrophobicity are presumed to be ideal to form stable emulsions [6], [7]. Hydrophobic bacteria possess an affinity for each other and undergo self-assembly at the oil-water interface, which enables them to resist coalescence and deformation [6]. Such bacteria thus have the potential to replace solid fat or synthetic surfactants and provide emulsions with an ‘all-natural’ designation [8].

Analysis of bacterial cell surface characteristics can accelerate the search for potential strains with high emulsion stabilizing properties. Microbial surface characteristics can be assessed by several methods, including microbial adhesion to hexadecane (MATH) [9] or solvents (MATS) [10]. Hexadecane is a nonpolar solvent that is devoid of electrostatic and hydrogen bond interactions, and hence microbial adhesion to this solvent can be used to deduce non-polar properties [11], and thereby surface hydrophobicity [10].

Bacteria are known to alter their surface compositions as a response to external influences including stressing conditions and chemical compounds, and various environmental factors. This property has been increasingly exploited to enhance surface hydrophobicity, creating promising Pickering particles for colloidal food materials[12]–[20]. A few naturally occurring bacteria and some yeast strains with inherent capabilities to serve as Pickering stabilizers have been previously described. For example, the formation of stable oil-in-water emulsion using natural cells of *Acinetobacter venetianus* RAG-1 and *Rhodococcus erythropolis* 20SE1-c have been demonstrated. These cells interacted with each other and formed surface films and emulsion gels, thereby hindering the coalescence of oil droplets. The changes in the interfacial tension was only negligible and the cells behaved as fine particles rather than synthetic surfactants [6]. Stable micron-sized Pickering-type colloidal particles could also be formed using cells of *S. cerevisiae*, *Lb. acidophilus* and *Strep. thermophilus*. These cells have been shown to adhere to the oil-water interface and provide long-term stability by preventing droplet coalescence and bulk phase separation [15].

Bacteria with hydrophobic surface characteristics have been considered rare in nature and hence most bacteria have been considered as unfit for Pickering stabilization by themselves due to the hydrophilic nature of their surfaces [6], [21]. The existence of intrinsic Pickering stabilization potential among different strains of bacteria seems to be evident based on some previous studies [6], [8], [15], [22], [23]. Nevertheless, comprehensive screening and investigation of large collections of lactic acid bacteria (LAB) strains, aimed at elucidating their intrinsic Pickering stabilization potential, have still been inadequate. At times, there can be variations in surface characteristics among strains of bacteria belonging to the same species and hence it could also be useful to consider similar properties in the search for strains with high emulsion stabilizing properties. Furthermore, the lack of consistent success to obtaining bacteria with enhanced surface hydrophobicity by applying such surface engineering approaches and challenges related to cost, labor, and perception towards modified organisms, emphasizes the need for essentially natural and complementary approaches for discovering attractive bacterial Pickering stabilizers.

Accordingly, the purpose of the present work is to overview the intrinsic Pickering stabilization potentials of bacterial strains belonging to different species within the former *Lactobacillus* genus by characterizing their cell surface hydrophobicity and charge. For selected strains, the correlation between these cell surface properties and Pickering emulsion stability was investigated.

## 2. Materials and Methods

### 2.1. Materials and chemicals

MRS broth and Agar Powder Bacteriological Grade were purchased from HiMedia (India), Hexadecane (density 0.77 g/mL) and KH_2_PO_4_ from Sigma-Aldrich (Steinheim, Germany) and Microtiter plates from Corning corporation (Corning, Germany). Medium-chain triglyceride (MCT) oil was obtained from AAK AB (Karlshamn, Sweden).

### 2.2. Bacterial strains

A total of 31 LAB strains belonging to 23 species of the former *Lactobacillus* genus were investigated in this study. Table 1 shows the former and current nomenclatures of the bacterial species and strains and the optimum temperatures used to cultivate them. All the strains were kindly obtained from strain collection of Department of Food Science, University of Copenhagen (Finn Kvist Vogensen, Personal communication). Unless otherwise indicated, strains were cultivated in MRS broth, using the indicated optimum temperatures (Table 1), and MRS agar was prepared by supplementing MRS broth with 2% (w/v) agar powder.

**Table 1.**
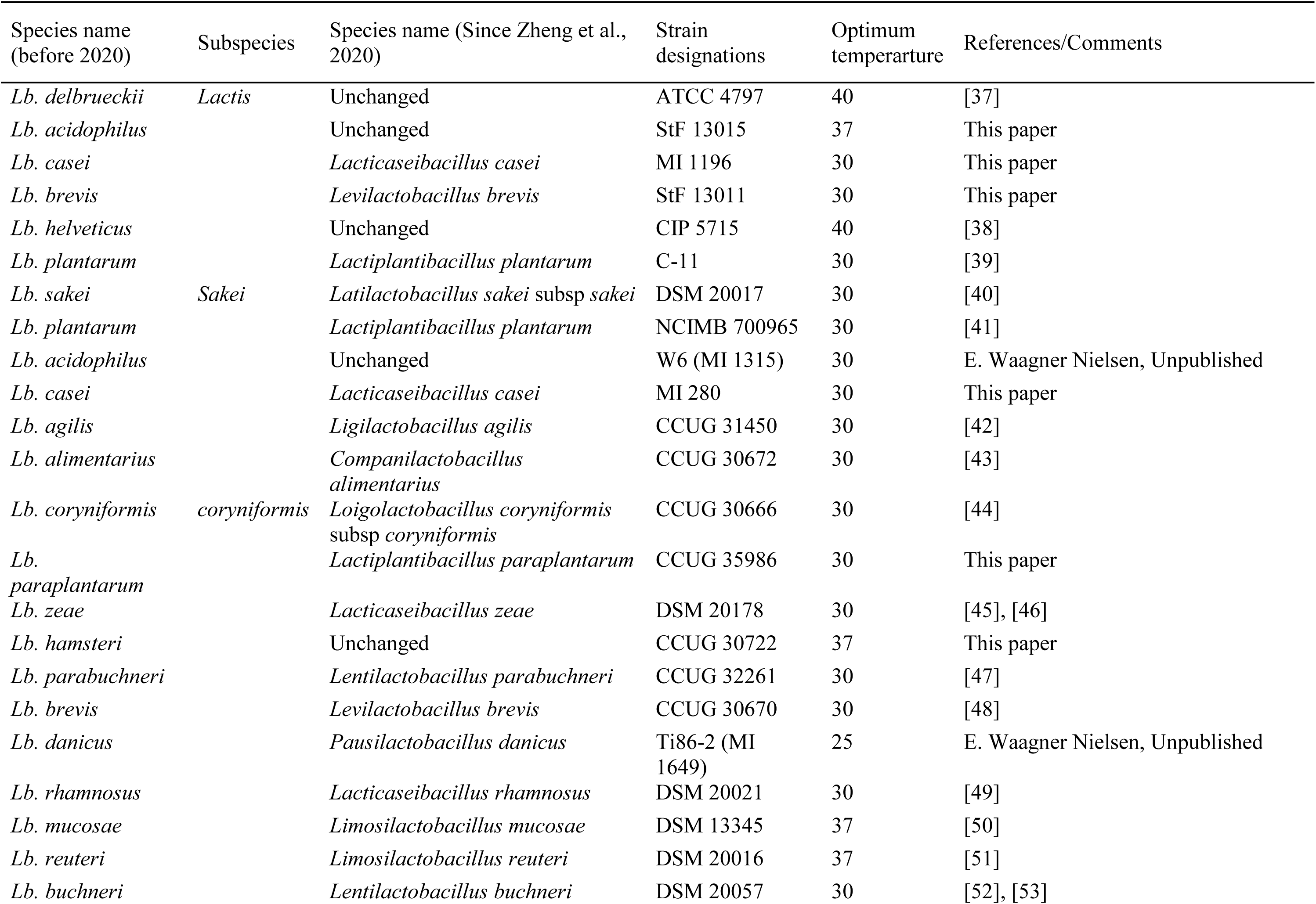
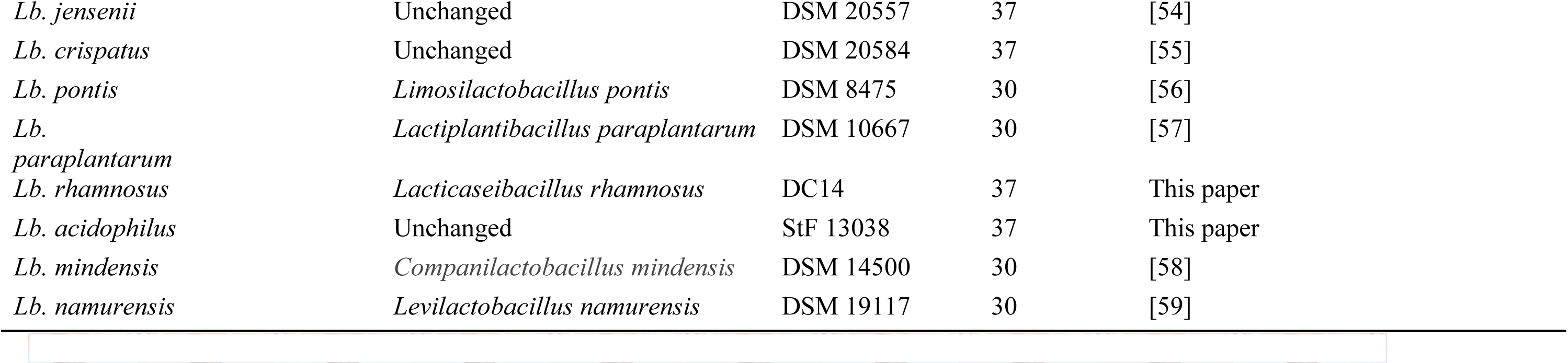
Summary of species strains of ex-*Lactobacillus* used in this study and corresponding optimum growth temperatures

| Species name<br>(before 2020) | Subspecies | Species name (Since Zheng et al.,<br>2020) | Strain<br>designations | Optimum<br>temperature | References/Comments |
| --- | --- | --- | --- | --- | --- |
| *Lb. delbrueckii* | *Lactis* | Unchanged | ATCC 4797 | 40 | [37] |
| *Lb. acidophilus* |  | Unchanged | StF 13015 | 37 | This paper |
| *Lb. casei* |  | *Lacticaseibacillus casei* | MI 1196 | 30 | This paper |
| *Lb. brevis* |  | *Levilactobacillus brevis* | StF 13011 | 30 | This paper |
| *Lb. helveticus* |  | Unchanged | CIP 5715 | 40 | [38] |
| *Lb. plantarum* |  | *Lactiplantibacillus plantarum* | C-11 | 30 | [39] |
| *Lb. sakei* | *Sakei* | *Latilactobacillus sakei* subsp *sakei* | DSM 20017 | 30 | [40] |
| *Lb. plantarum* |  | *Lactiplantibacillus plantarum* | NCIMB 700965 | 30 | [41] |
| *Lb. acidophilus* |  | Unchanged | W6 (MI 1315) | 30 | E. Waagner Nielsen, Unpublished |
| *Lb. casei* |  | *Lacticaseibacillus casei* | MI 280 | 30 | This paper |
| *Lb. agilis* |  | *Ligilactobacillus agilis* | CCUG 31450 | 30 | [42] |
| *Lb. alimentarius* |  | *Companilactobacillus*<br>*alimentarius* | CCUG 30672 | 30 | [43] |
| *Lb. coryniformis* | *coryniformis* | *Loigolactobacillus coryniformis*<br>subsp *coryniformis* | CCUG 30666 | 30 | [44] |
| *Lb.*<br>*paraplantarum* |  | *Lactiplantibacillus paraplantarum* | CCUG 35986 | 30 | This paper |
| *Lb. zeae* |  | *Lacticaseibacillus zeae* | DSM 20178 | 30 | [45], [46] |
| *Lb. hamsteri* |  | Unchanged | CCUG 30722 | 37 | This paper |
| *Lb. parabuchneri* |  | *Lentilactobacillus parabuchneri* | CCUG 32261 | 30 | [47] |
| *Lb. brevis* |  | *Levilactobacillus brevis* | CCUG 30670 | 30 | [48] |
| *Lb. danicus* |  | *Pausilactobacillus danicus* | Ti86-2 (MI<br>1649) | 25 | E. Waagner Nielsen, Unpublished |
| *Lb. rhamnosus* |  | *Lacticaseibacillus rhamnosus* | DSM 20021 | 30 | [49] |
| *Lb. mucosae* |  | *Limosilactobacillus mucosae* | DSM 13345 | 37 | [50] |
| *Lb. reuteri* |  | *Limosilactobacillus reuteri* | DSM 20016 | 37 | [51] |
| *Lb. buchneri* |  | *Lentilactobacillus buchneri* | DSM 20057 | 30 | [52], [53] |

|  |  |  |  |  |
| --- | --- | --- | --- | --- |
| *Lb. jensenii* | Unchanged | DSM 20557 | 37 | [54] |
| *Lb. crispatus* | Unchanged | DSM 20584 | 37 | [55] |
| *Lb. pontis* | *Limosilactobacillus pontis* | DSM 8475 | 30 | [56] |
| *Lb. paraplantarum* | *Lactiplantibacillus paraplantarum* | DSM 10667 | 30 | [57] |
| *Lb. rhamnosus* | *Lacticaseibacillus rhamnosus* | DC14 | 37 | This paper |
| *Lb. acidophilus* | Unchanged | StF 13038 | 37 | This paper |
| *Lb. mindensis* | *Companilactobacillus mindensis* | DSM 14500 | 30 | [58] |
| *Lb. namurensis* | *Levilactobacillus namurensis* | DSM 19117 | 30 | [59] |

### 2.3. Preparation of bacterial cells

A small portion of the stock culture was streaked onto an appropriate agar plate and incubated overnight with the optimum growth conditions. A well-separated colony was picked from the plate and transferred into 10 mL broth and incubated overnight at the optimum growth condition and in the absence of shaking. For assessment of zeta potential and cell surface hydrophobicity, cells were harvested by centrifugation at 4000 x g for 15 min at 4°C, washed twice by resuspending with sterile Milli-Q water (Merck Millipore) and collected by centrifugation at 5000 x g for 15 min at 4°C. For preparation and analysis of Pickering emulsions, 250 µL of the overnight culture was further inoculated into 100 mL broth, incubated overnight at appropriate conditions without shaking, and cells were collected as described above. Cells were resuspended in PBS buffer pH 7.4 (1 mL for those for cell surface hydrophobicity and zeta potential analysis, and 3 mL for Pickering emulsion analysis), and kept in the refrigerator until analysis (maximum for a week).

### 2.4. Measurement of cell surface hydrophobicity

Cell surface hydrophobicity was assessed using MATH as previously described [10], [24]. In brief, a portion of the bacterial cell suspension was transferred into Eppendorf tube and centrifuged at 10,000 x g. The supernatant was discarded, and the pellet resuspended with 1 mL 10 mM KH_2_PO_4_ solution pH 4.0. The absorbance of the cell suspension was adjusted to OD_600_ of 0.5±0.1 (A0) (by measuring 150 µL sample in a microtiter plate using Varioscan (ThermoScientific). Two hundred fifty (250) µL of the OD-adjusted suspension was mixed with 45 µL hexadecane in a microtiter plate and the mixture thoroughly mixed by pipetting up and down (90 times). The mixture was let to stand for 15-30 min at room temperature to allow complete phase separation. Two hundred (200) µL of the aqueous phase was transferred onto microtiter plate. From the first transfer, 150 µL was transferred to a new microtiter plate and OD_600_ was measured (A1). The percentage of MATH was calculated using the following equation:

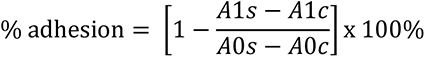

Where A1s and A1c are the corresponding absorbances of the sample and control after cell adhesion to hexadecane, and A0s and A0c are the corresponding absorbances of the sample and control prior to the assay. All the results were obtained from triplicated measurements and data are presented as average ± standard deviation.

### 2.5. Measurement of zeta potential

A small portion of the bacterial cell suspension (2.5 µL in PBS buffer pH 7.4) was transferred into Eppendorf tube containing 1 mL sterile Milli-Q water and centrifuged at 10,000 x g. The supernatant was discarded, and the pellet resuspended with sterile 1 mL Milli-Q water. Zeta potential (ZP) was measured using Zeta Sizer nano ZSP (Malvern Analytical) at 25°C. Results were presented as average ± standard deviation of duplicate measurements, each of which were measured as technical triplicates.

### 2.6. Preparation and analysis of Pickering emulsions

Emulsion experiments were carried out essentially as previously described [25] on a total of 12 strains. Detailed description of the bacterial strains selected for emulsion studies is provided in Table S2 (Supplementary Material 1). In brief, a portion of the bacterial cell suspension in PBS (pH 7.4) was transferred into 15 mL centrifuge tubes (with volume indicators) and pelleted at 4,000 x g for 15 min. The supernatant was discarded, and the pellet resuspended with 5 mL 10 mM KH_2_PO_4_ solution (pH 4.0). The optical density (OD_600_) was adjusted to ca. 2.5 (for measurement accuracy, 10-fold diluted cell suspension was used). A 150 µL sample was used for OD measurement. OD was measured using Varioscan using clear microtiter plates. Five (5) ml of the OD-adjusted sample was mixed with equal volume of MCT oil, and the sample mixture was thoroughly mixed using a high-shear mixer (T25 digital Ultra-Turrax®, IKA) at 24000 rpm for 60 s. Each sample was analyzed in duplicate.

Prepared emulsions were stored at room temperature and visual stability was analyzed after 1, 2 and 3 weeks. The volumes of the emulsified material and that of the entire sample preparation were noted during each measurement timepoint alongside storage. These volumes were used to calculate the emulsions’ relative volume, defined as the ratio in percentage of the volume of the emulsified material relative to the entire volume of the sample preparation. Visual stability of the emulsions was eventually estimated by assessing the shift in emulsion relative volume over time.

Emulsions were further analyzed using a laser diffraction particle size analyzer (Mastersizer 3000, Malvern Instruments, Workshire, UK) at 25°C on day 0. Emulsion stability was investigated by the time-dependent shifts in (i) the mean droplet diameter over volume (aka equivalent volume mean or d(4,3)), (ii) droplet size distribution (DSD, emulsion droplet diameters at 10%, 50%, and 90% cumulative volumes [Dv10, Dv50 and Dv90, respectively]) and (iii) relative span (i.e., the width of the distribution of droplet sizes [26]). Before each measurement, samples were gently flipped up and down twice to re-disperse the creaming droplets into the aqueous phase to avoid multiple scattering effects. Additionally, as the emulsion produced with strain *Lb. acidophilus* StF 13015 had extremely firm consistency, and as tube inversion alone was not sufficient to re-disperse the droplets into the aqueous phase, more vigorous force was applied to this sample. Samples were then slowly added into the sample dispersion unit containing Milli-Q water and mixed at a stirring velocity of 2200 rpm before they were pumped into the optical chamber. The obscuration range was set between 2% and 15%. The refractive and absorption indices were set to 1.47 and 0.01, respectively. The following equation was used to calculate relative span.

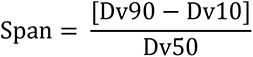

Where, Dv10, Dv50 and Dv90 are the maximum droplet diameters, below which 10%, 50% and 90% of the volume of emulsion exists, respectively [27]. That is, [Dv90-Dv10] is the range of the data and Dv50 the median diameter [28].

## 3. Results

### 3.1. Zeta Potential of Strains

The occurrence of bacteria with specific zeta potential values is summarized (Figure S1b, Supplementary Material 2). The mean zeta potential values measured for each individual strains are also provided in Table S1 (Supplementary Material 1). On the frequency plot, the strains appeared to be roughly divided into two discrete groups. One of these groups comprised of strains with highly negative zeta potential and appeared to form a major peak on the distribution plot that spanned from -43.76 to -19.23 mV. This group constituted most of the tested strains, i.e., ∼77.42% (24 out of 31). The mean and median zeta potential values were -32.13±5.62 mV and -32.01 mV, respectively. The other group comprised of cells with relatively neutral zeta potentials and tended to form a minor peak on the distribution plot that spanned from -12.51 mV to +1.14 mV. This group constituted a total of 7 strains (i.e., ∼22.58% of the tested strains), including *Lacticaseibacillus casei* MI 1196 and *Limosilactobacillus pontis* DSM 8475 -strains with exclusively positive zeta potentials (+1.14±0.46 mV and +0.21±0.80 mV, respectively). The strains constituting the minor group had mean and median zeta potential values of -4.75±5.04 mV and -3.96 mV, respectively. In general, except from a few of the tested strains, the zeta potential values were all negative.

### 3.2. Surface Hydrophobicity of Strains

We determined surface hydrophobicity using MATH, i.e., by measuring the extent of cell surface adhesion to hexadecane. Figure S1a (Supplementary Material 2) summarizes the distribution of strains according to their MATH values. The mean MATH values measured for all individual strains are also provided in Table A1 (Supplementary Material 1). The tested strains predominantly demonstrated certain degree of hydrophobicity (MATH > 30%), while the rest exhibited a low degree hydrophobicity (MATH < 30%). The fraction of strains that exhibited a certain degree of hydrophobicity (i.e., MATH > 30%) was ∼58% (18 out of 31). Of these, 6 strains were of weak hydrophobicity (30% < MATH < 50%), 5 strains moderate hydrophobicity (50% < MATH < 75%), and 7 strains demonstrated high hydrophobicity (MATH > 75%). The ratio of strains that exhibited low hydrophobicity (i.e., MATH < 30%) was ∼42% (13 out of 31) and these strains were referred to as ‘highly hydrophilic’.

### 3.3. Correlation Between Zeta Potential and Surface Hydrophobicity

Since there was considerable variation in zeta potential and surface hydrophobicity among the tested strains, the presence of correlation was assessed by plotting the results of zeta potential and surface hydrophobicity of all individual strains against each other (Figure 1). The finding demonstrated a complex relationship between zeta potential and surface hydrophobicity, thus ruling out the possibility of strong linear correlation between these measurements (R^2^=0.0198).

**Figure 1.**
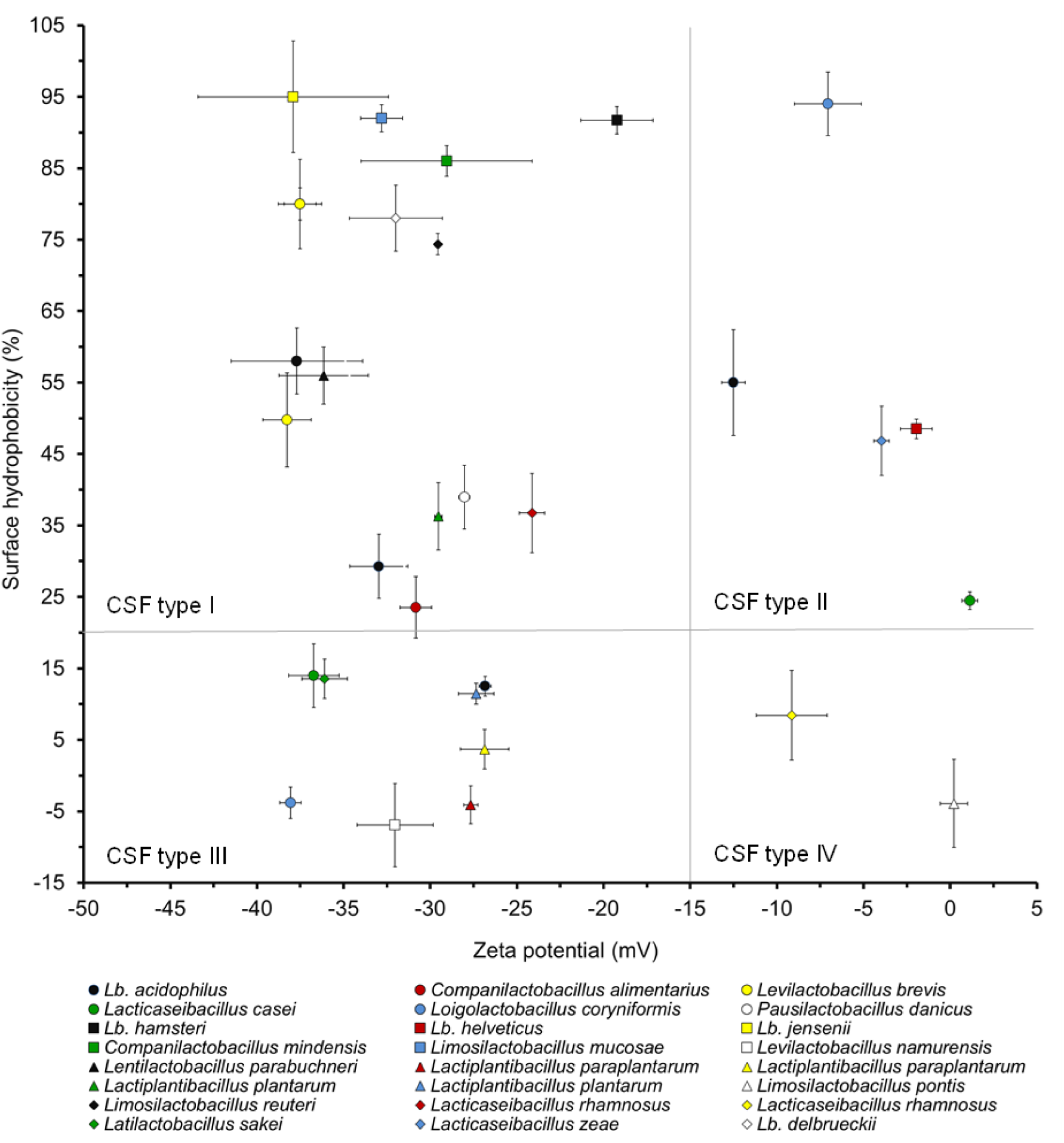
Coordinates of zeta potential vs. surface hydrophobicity highlighting 4 major cell surface features (CSFs) among 31 ex-*Lactobacillus* strains belonging to 24 species. The cell surfaces of both type I and II strains were highly hydrophobic, whereas the zeta potentials varied from highly negative (< -15 mV, type I) to relatively neutral (> - 15 mV, type II). The same applied to type III and IV strains, both of which had less hydrophobic cell surfaces, but highly negative (type III) or relatively neutral (type IV) zeta potentials.

The distribution of strains on the zeta potential vs. hydrophobicity plot highlighted four major cell surface features (CSF) amongst the strains (Figure 1), designated here as type I, II, III and IV.

Strains with CSF type I were characterized by having certainly negative zeta potential (-43.76 mV ≤ zeta potential ≤ -19.23 mV) and high hydrophobicity (MATH >30%). These strains included *Lb. acidophilus* StF 13015, *Levilactobacillus brevis* StF 13011, *Levilactobacillus brevis* CCUG 30670, *Lentilactobacillus buchneri* DSM 20057, *Pausilactobacillus danicus* Ti86-2, *Lb. delbrueckii* subsp. *lactis* ATCC 4797, *Lb. hamsteri* CCUG 30722, *Lb. jensenii* DSM 20557, *Companilactobacillus mindensis* DSM 14500, *Limosilactobacillus mucosae* DSM 13345, *Lentilactobacillus parabuchneri* CCUG 32261, *Lactiplantibacillus plantarum* NCIMB 700965, *Limosilactobacillus reuteri* DSM 20016 and *Lacticaseibacillus rhamnosus* DC14, thereby representing the most dominant and diverse, representing about 45% of the tested strains (14 out of 31 strains). Those strains with CSF type II, namely *Lb. acidophilus* StF 13038, *Lb. crispatus* DSM 20584, *Lb. helveticus* CIP 5715,

*Lacticaseibacillus zeae* DSM 20178, tend to have relatively neutral zeta potential (- 12.51 mV ≤ zeta potential ≤ +1.14 mV) but coupled with high hydrophobicity (MATH > 30%) and represented about 13% of the tested strains (4 out of 31).

Strains designated as CSF type III, namely *Lb. acidophilus* W6, *Ligilactobacillus agilis* CCUG 31450, *Companilactobacillus alimentarius* CCUG 30672, *Lacticaseibacillus casei* MI 280, *Loigolactobacillus coryniformis subsp. coryniformis* CCUG 30666, *Levilactobacillus namurensis* DSM 19117*, Lactiplantibacillus paraplantarum* CCUG 35986, *Lactiplantibacillus paraplantarum* DSM 10667, *Lactiplantibacillus plantarum* C-11 and *Latilactobacillus sakei subsp. sakei* DSM 20017, defined by certainly negative zeta potential (-43.76 mV ≤ zeta potential ≤ -19.23 mV) and low hydrophobicity (MATH <30%), represented about 32% of the strains analyzed (10 out of 31 strains). Cells with CSF type IV, namely *Lacticaseibacillus casei* MI 1196, *Limosilactobacillus pontis* DSM 8475 and *Lacticaseibacillus rhamnosus* DSM 20021, comprised of relatively neutral zeta potential (-12.51 mV ≤ zeta potential ≤ +1.14 mV) combined with low hydrophobicity (MATH <30%). Strains with this latter CSF type appeared to be the least frequent (i.e., ∼10% [only 3 out of 31 strains]).

### 3.4. Effect of Selected Strains on Emulsion Formation and Stability

To analyze whether incorporation of cells with high surface hydrophobicity will have an impact on emulsion formation and stability, emulsions were Pickering-stabilized with CSF type I strains (see Table A1), and the relative changes in the macroscopic appearance of the Pickering emulsions, as well as the shift in D-values, relative span and d(4,3) at various storage times (0, 1, 2 and 3 weeks) were analyzed.

#### 3.4.1. Macroscopic appearances of the emulsions

##### 3.4.1.1. Relative emulsion volume

For most of the emulsions, it took generally a few days to separate into different phases, and immediate phase separation was rarely observed. Macroscopic appearances of representative emulsions are presented (Figure S2, Supplementary Material 2). To estimate the physical stability of the emulsions, the relative volume of the emulsified material at various storage times was determined. The findings for the first and subsequent weeks of storage are presented (Figure 3). The mean relative volumes analyzed for individual emulsions following storage for a week are shown (Table S2, Supplementary Material 1).

Note that the initial ratio between the aqueous and organic phase volumes was 1:1. Nonetheless, the relative volume of the emulsified material turned out to be more than half of the total sample volume for emulsions of *Lb. acidophilus* StF 13015, *Limosilactobacillus mucosae* DSM 13345, *Lb. jensenii* DSM 20557 and *Lb. crispatus* DSM 20584 (Figure 3). For these emulsions, the relative volume of the none-emulsified sample fraction was thus decreased to below 50%. Over the course of storage of these emulsions, no major reduction in relative emulsion volume was observed, except for *Lb. crispatus* DSM 20584 that showed an overall reduction of ∼13% (from ∼57% to ∼44%, slope=-0.85±0.00) following storage for three weeks (Figure 3). For the remaining emulsions, the relative emulsion volume mimicked that of the initial organic phase volume, i.e., roughly 50% (Figure 3 and Table S2, Supplementary Material 2). Most of these emulsions did not exhibit major relative emulsion volume reduction upon storage, except *Lb. hamsteri* CCUG 30722 and *Companilactobacillus mindensis* DSM 14500, both of which showed a total reduction of ∼12% following storage for three weeks (Figure 3). Thus, most strains are capable of maintaining the stability of the forged emulsion and, considering the limted number of strains with natural propnesity to forge massive emulsion, there is seeminlgy little inter-dependence between these two natural strain capabilities.

##### 3.4.1.2. Consistency and cell precipitation

The majority of the emulsions had a consistency that, following a couple of gentle tube inversions prior to analysis, resulted in dispersion of the creaming emulsion droplets into the underlying aqueous solution. This was, however, not the case for *Lb. acidophilus* StF 13015 emulsion, which appeared to have very fine gross texture and extremely firm consistency (data not shown). As a result, the tube had to be more vigorously agitated to disperse the emulsified material into the aqueous solution, suggesting that such firm texture was the result of more robust adsorption of cells onto the interface, thereby enhancing stability of the emulsion.

A common phenomenon related to storage of the emulsions was cell precipitation. For *Lb. crispatus* DSM 20584 and *Lb. hamsteri* CCUG 30722, cell precipitation was observed relatively early (as early as one day after preparation). For many of the strains, cell precipitation increased steadily alongside storage, with *Lb. crispatus* DSM 20584 and *Lb. hamsteri* CCUG 30722 showing the highest increase, while *Lb. acidophilus* StF 13015, *Levilactobacillus brevis* CCUG 30670 and *Companilactobacillus mindensis* DSM 14500 exhibiting the lowest increase (data not shown). This proves that cells of some ex-*Lactobacillus* strains have natural propensity to undergo rapid precipitation, and this could have negative impact on adsorption of cells onto the interface, thereby affecting emulsion stability.

#### 3.4.2. Size distribution of emulsion droplets

Droplet size distribution (DSD) plots of individual emulsions were analyzed to highlight the dominant droplet sizes. DSD plots for all the emulsions are shown (Figure S3, Supplementary Material 2). Virtually, all DSD plots demonstrated minor and major peaks, the intensity of which had occasional variations among the different strains of ex-*Lactobacillus*. The minor and major peaks spanned from 1-25 µm and from 30-200 µm size ranges, respectively. The highest value for the minor peaks was typically ∼13 µm for majority of the emulsions. For the major peaks, the highest value varied roughly between 80 and 90 µm. Importantly, the major peak of *Lb. acidophilus* StF 13015 spanned from 15 µm to 200 µm, with the curve topping at around 62 µm. A minor peak was less evident in this emulsion (Figure S3, Supplementary Material 2), possibly suggesting notable connection between the underlying elements and overall emulsion droplet size.

DSD plots generated at different timepoints were used to explore emulsion stability. Stable emulsions were defined by DSD plots of high overall uniformity over the whole duration of storage, whereas with the less stable emulsions, the DSD plots varied with storage. Examples of DSD plots of stable and less-stable emulsions are presented (Figure 2A and 2B, respectively). It was found that *Lb. hamsteri* CCUG 30722, for instance, provided nearly uniform DSD plots regardless of storage for up to three weeks at room temperature (Figure 2A). In general, DSD plots for *Lb. delbrueckii lactis* DSM 20076, *Lacticaseibacillus casei* MI 280, *Lb. hamsteri* CCUG 30722, *Lentilactobacillus parabuchneri* CCUG 32261, *Limosilactobacillus mucosae* DSM 13345, *Lb. jensenii* DSM 20557 and *Lb. crispatus* DSM 20584 retained uniformity despite continuous storage and measurement (Figure A2). Contrary to these, plots for *Lb. acidophilus* StF 13015, *Levilactobacillus brevis* CCUG 30670, *Lentilactobacillus buchneri* DSM 20057, *Lb. acidophilus* StF 13038 and *Companilactobacillus mindensis* DSM 14500 showed a more variable trend in response to storage (Figure S3). The variaition in density during storage appeared to be negligible for smaller droplets, indicating the positive influence of such droplets for the overall stability of the emulsions. Meaningful differences could be observed in the fraction of larger droplets in response to storage, thereby signifying a more destabilizing role (Figure S3).

**Figure 2.**
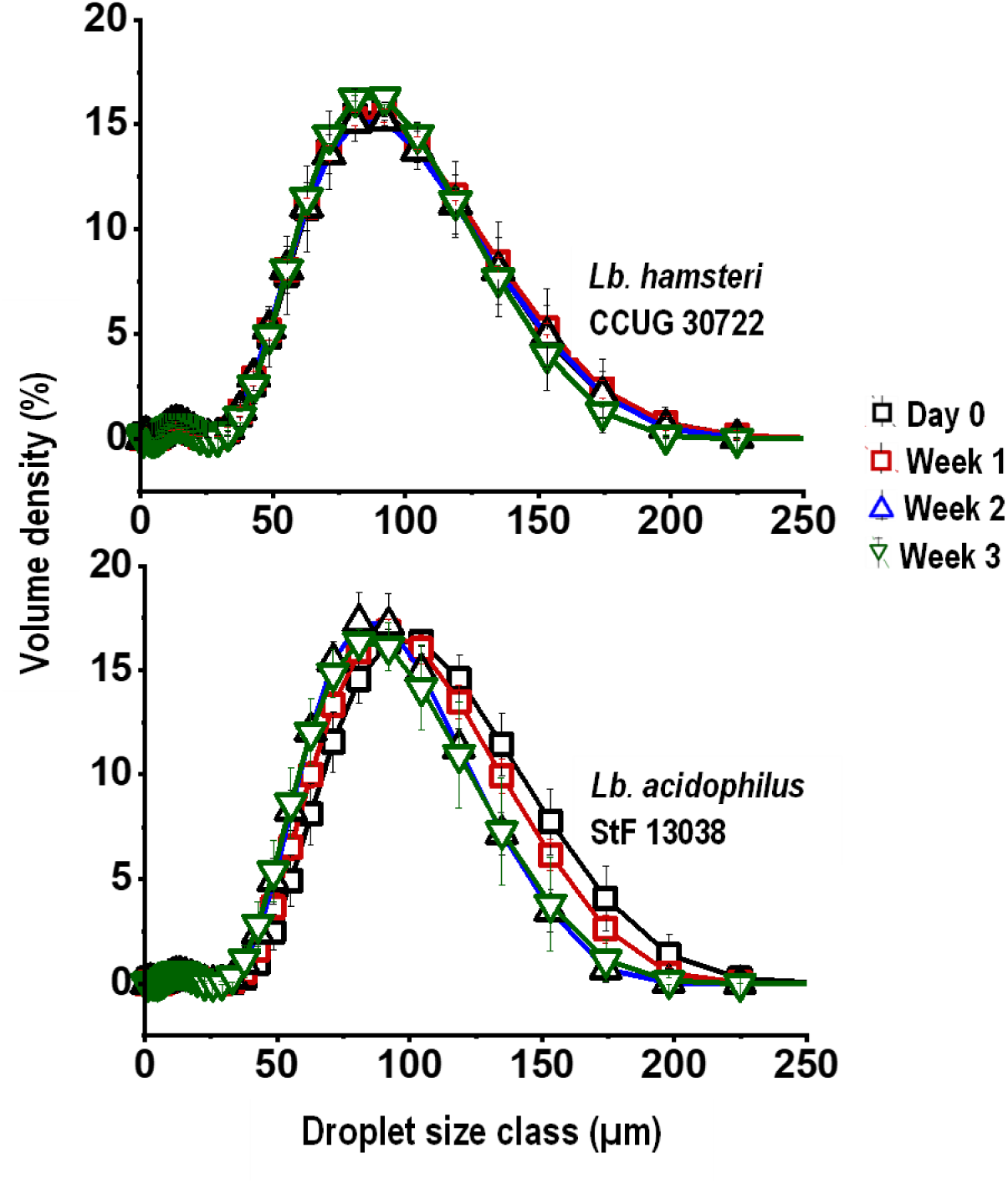
Examples of emulsion DSD plots of *Lb. hamsteri* CCUG 30722 and *Lb. acidophilus* StF 13038 generated at the day of preparation and susequent weeks, demonstrating relatively stable and less stable emulsions, respectively. Minor peaks at around 10 µm of the *Lb. hamsteri* CCUG 30722 plot correspond to bacterial cells not adsorbed to the interface. Emulsions were prepared and measured in duplicates, and samples were gently inverted twice prior to measurement.

The high relative stability in density of smaller droplets over time could also be reflected by corresponding Dv10 measurements (Figure S4, Supplementary Material 2). The lowest and highest Dv10 values were measured to be 30.05±14.45 µm and 61.11±2.20 µm. Two other parameters, namely Dv50 and Dv90, had higher values and showed larger differences. The lowest and highest values were measured to be 54.53±25.46 µm and 95.42±3.75 µm for Dv50 and 100.78±10.68 µm and 146.50±6.94 µm for Dv90. Although all lowest D-values were exclusively coupled with *Lb. acidophilus* StF 13015, the highest values were associated with *Limosilactobacillus mucosae* DSM 13345 (Dv10 and Dv50, both on day 0) and *Lb. acidophilus* StF 13038 (Dv90, day 0). Regardless of these differences, all the three parameters were less significantly correlated with surface adherence to hexadecane (Figures S5, Supplementary Material 2) and surface charge (Figures S6, Supplementary Material 2). The same trends were found for the relative spans, defined as the width of droplet size distribution (Figure S7, Supplementary Material 2). Like the D-values, this parameter was also less significantly correlated with surface adherence to hexadecane and surface charge (Figure S5 and S6, Supplementary Material 2).

Further analysis of the relative span (Figure S7, Supplementary Material 2) revealed the width of the droplet size distributions of all emulsions, except of *Lb. acidophilus* StF 13015 (*span*=1.37±0.18), to be initially narrow (*Span*=0.91±0.02). *Limosilactobacillus mucosae* DSM 13345 was the most efficient in terms of providing the lowest span, i.e., 0.88±0.03. Over the course of storage, the relative span varied only slightly (mean span=1.97±1.82, 1.06±0.18, 0.98±0.12 and 0.98±0.11 for day 0, week 1, 2 and 3, respectively). It was only *Levilactobacillus brevis* CCUG 30670 that showed a large variation in relative span between sample replicates, i.e., it exhibited large standard deviations that persisted throughout the entire duration of storage. The relative span of all the other strains was rather stable, ruling out the possibility of major drift in the size distribution of smallest and largest droplets.

Equivalent volume mean or d(4,3) analysis was employed to determine the size of droplets that constitute the bulk of the sample volume (Figure 3). Analysis of this parameter at various storage times was employed to determine droplet storage variability. At the day of preparation, the d(4,3) measurement averaged 88.33±9.48 µm. Here, the lowest and highest values corresponded to *Lb. acidophilus* StF 13015 (61.42±25.67 µm) and *Limosilactobacillus mucosae* DSM13345 (99.49±4.01 µm), respectively. After storage for a week, the mean value slightly decreased (85.50±6.71 µm). At this storage time, the lowest and highest values corresponded to *Lb. acidophilus* StF 13015 (67.73±3.21 µm) and *Lb. acidophilus* StF 13038 (93.27±2.56 µm). After two weeks of storage, the mean value slightly decreased (83.78±8.62 µm). The lowest and highest values corresponded to *Lb. acidophilus* StF 13015 (62.35±8.37 µm) and *Limosilactobacillus mucosae* DSM13345 (96.04±1.32), respectively. After three weeks of storage, the mean value remained nearly the same as that determined at the second week (82.28±7.49 µm). Although *Lb. acidophilus* StF 13015 had still the lowest value (62.13±10.61 µm), it was *Companilactobacillus mindensis* DSM 14500 that showed the highest value (92.25±4.64 µm). It is therefore *Lb. acidophilus* StF 13015 that generally offered smaller emulsion droplets as compared to the rest of the strains (Figure 3).

Figure 3 also shows the storage variability of d(4,3) values of all individual emulsions. Accordingly, five of the emulsions (namely *Lentilactobacillus buchneri* DSM 20057, *Lb. hamsteri* CCUG 30722, *Lb. crispatus* DSM 20584*, Lb. acidophilus* StF 13015 and *Companilactobacillus mindensis* DSM 14500) had low d(4,3) overall storage variability. An equally high number of emulsions (i.e., *Limosilactobacillus mucosae* DSM 13345, *Lb. jensenii* DSM 20557, *Levilactobacillus brevis* CCUG 30670, *Lentilactobacillus parabuchneri* CCUG 32261 and *Lb. delbrueckii* subsp. *lactis* DSM 20076) exhibited moderate d(4,3) storage variability. In these latter emulsions, an overall reduction in droplet size over time was detected (Figure 3). For the rest the emulsions (namely *Lacticaseibacillus casei* MI 280 and *Lb. acidophilus* StF 13038), a high d(4,3) variability was observed during storage. In general, storage of the cell- stabilized emulsions was accompanied by an overall decrease in droplet size (Figure 3).

**Figure 3.**
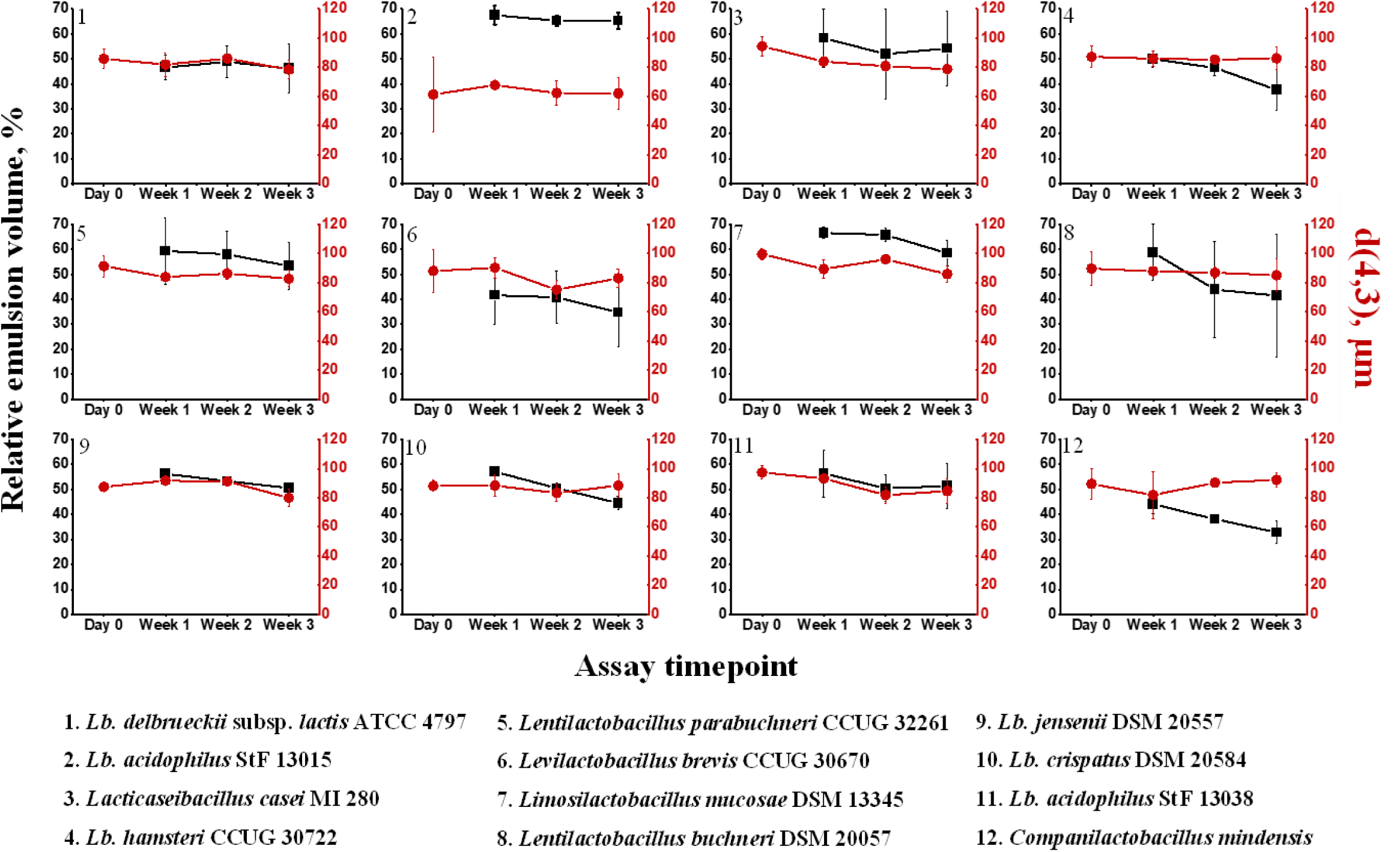
Relative volumes (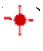) and equivalent volume means (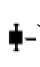) of ex-*Lactobacillus* strains-stabilized emulsions measured at day 0, week 1, 2 and 3. The relative volume is the emulsified material’s volume relative to the total sample volume, expressed in percentage terms. Experiments were carried out in duplicates and data are presented as average ± standard deviation.

## 4. Discussion

Due to their GRAS (generally recognized as safe) status, in addition to their centuries-long use in the manufacture of dairy and fermented food products, strains of the ex- *Lactobacillus* species have been increasingly considered for Pickering emulsion stabilization [6], [13], [15]–[20]. However, only a limited number of strains have as yet been investigated as potential Pickering particles, and it is mainly surface-modified cells that have been studied. In this study, the intrinsic Pickering emulsion formation and stabilization abilities of 12 strains belonging to 11 species of the ex-*Lactobacillus* was evaluated, following prior analysis of the cell surfaces properties of 31 strains belonging to 24 species.

One of the most common surface characteristics of cells is their adhesion to solvents, which can be used to determine the non-polar properties of bacteria [11]. Partitioning studies often employ hexadecane as a solvent and such organic phase typically occupies the upper phase due to a lower density (0.77 vs. 0.99 g/mL for water). The partitioning of bacterial cells is affected by gravity/turbulence [17]. Thermal motion alone would not be sufficient to keep Gram-positive bacteria, with densities of ∼1.2 g/mL [29], suspended in hexadecane and water for long time [17], thus resulting in eventual cell sedimentation. Thus, reliable analysis of surface hydrophobicity of cells by the MATH assay necessitated prior evaluation of a suitable duration of incubation that provides minimal loss of cells by sedimentation. As part of preliminary survey, we observed that two of the strains, namely *Lb. hamsteri* CCUG 30722 and *Lb. crispatus* DSM 20584, were prone to relatively fast sedimentation, where visible sedimentation was observable after ∼45 min of standing. This is the main reason for allowing a waiting time of only 15-30 min for the MATH assay, thus minimizing the impact of cell sedimentation on the overall analysis.

Previous studies have suggested that most ex-*Lactobacillus* cell surfaces were intrinsically hydrophilic [6], [21], [24], [30]. Importantly, the present finding is in contrast to such previous claims, as a relatively large proportion (∼58%) of strains with high hydrophobicity (MATH >30%) is found. This screening of relatively large proportion of ex-*Lactobacillus* strains with high hydrophobicity thus clearly demonstrate the existence of naturally occurring, hydrophobic strains in the population.

Our result strongly indicated the absence of linear correlation between zeta potential and surface hydrophobicity. Similar results were previously reported [31]. Bacterial colloidal properties could be strongly influenced by cell-wall heterogeneities and classical colloidal concepts, like electrostatic potential and hydrophobicity, alone may not be sufficient to describe the complexity behind bacterial surface properties [32]. Structural organization of the various components and the conformational degrees of freedom of the polymeric surface constituents also play important roles [32]. The propensity of a bacterium to adhere to surfaces and to bind to polymeric constituents is determined by the global physicochemical nature of the outer layer of the cell wall, the conformation of the surface macromolecules and the susceptibility of the surface toward external perturbations [32]. Thus, the lack of linear correlation between zeta potential and surface hydrophobicity in this and other studies [31] strongly emphasizes the complex nature of the cell surface characteristics of the tested ex-*Lactobacillus* strains.

Nitrogen-rich constituents on the surfaces of ex-*Lactobacillus* cells have major positive influence on surface hydrophobicity [33]. Lipoteichoic acids, on the other hand, have been shown to impart negatively charged and, at the same time, hydrophobic surfaces to ex-*Lactobacillus* strains [32]. It was interesting to observe the latter phenomenon in almost half of our strains, namely those with CSF type I, which appeared to have negative zeta potential and at the same time high surface hydrophobicity. In one of the other CSF types, namely type III, the influence of negative zeta potential on surface hydrophobicity was rather inverse. As CSF type II represented only about a third of the tested strains, compared to type I, which represented almost half of the tested strains, we tentatively suggest that the abundance of negative charge on the surfaces of ex- *Lactobacillus* cells often has positive influence on cell surface hydrophobicity. We suggest also that cell surface neutrality generally has a lesser influence on surface hydrophobicity, as observed for CSF type II (minor positive impact) or IV (minor negative impact) strains. The weakly charged and hydrophilic surface characteristics of those strains with SCF type IV could be due to abundance of polysaccharides, which, according to previous reports [32], are generally weakly charged and hydrophilic.

Our emulsion data strongly suggested that only four strains, namely *Lb. acidophilus* StF 13015, *Limosilactobacillus mucosae* DSM 13345, *Lb. crispatus* DSM 20584 and *Lb. jensenii* DSM 20557, were capable of imparting higher ratio of the emulsified sample fraction, emphasizing that these four strains have natural abilities to form large Pickering emulsion volumes. Despite this difference in emulsion volume among the different strains of ex-*Lactobacillus*, the ratio of the emulsified sample fraction was generally stable during storage in all the emulsions, except for *Lb. crispatus* DSM 20584, *Lb. hamsteri* CCUG 30722 and *Companilactobacillus mindensis* DSM 14500. For *Lb. crispatus* DSM 20584 and *Lb. hamsteri* CCUG 30722, gradual loss of cells over time could explain the observed loss of macroscopic emulsion stability during storage, as these strains typically had rapid natural sedimentation. For *Companilactobacillus mindensis* DSM 14500, a strain with less rapid natural sedimentation, gradual loss of macroscopic emulsion stability seems to be a more dynamic phenomenon, involving events other than loss of cells.

Previous studies have reported that emulsions stabilized by ex-*Lactobacillus* strains had droplet sizes in the range of 30-300 [15] or 30-120 µm [17]. The droplet sizes of the emulsions stabilized by the strains in this study had major peaks over size range of 30- 200 µm, thus presenting remarkable agreement with the previous findings [15], [17]. Based on consistency of DSD curves of various timepoints, emulsions of *Lb. delbrueckii lactis* DSM 20076, *Lacticaseibacillus casei* MI 280, *Lb. hamsteri* CCUG 30722, *Lentilactobacillus parabuchneri* CCUG 32261, *Limosilactobacillus mucosae* DSM 13345, *Lb. jensenii* DSM 20557 and *Lb. crispatus* DSM 20584 were presumed to be stable. More frequently, the size distribution of larger droplets appeared to be less consistent over time, signifying that these droplets were more dynamically affected by emulsion destabilizing events. In contrast, the size distribution of smaller droplets appeared to be adequately consistent despite of continuous storage, thereby contributing profoundly for the overall stability of the emulsions.

The storage stability of majority of the emulsions was highly promising based on Dv10 and Dv50 measurements, except for *Companilactobacillus mindensis* DSM 14500, which showed less stable Dv10. Consistent with the notion that larger emulsion droplets are often prone to emulsion destabilizing processes, such as coalescence and flocculation [34], [35], the majority of the emulsions produced in this study were relatively less stable at 90% cumulative volumes. Accordingly, only emulsions of *Lb. hamsteri* CCUG 30722*, Lentilactobacillus buchneri* DSM 20057 and *Companilactobacillus mindensis* DSM 14500 had stable Dv90 measurements. This indicated that the observed variation in DSD was predominantly linked to Dv90. Observation of a drastic decline in Dv90 in emulsions of *Lacticaseibacillus casei* MI 280 and *Lb. acidophilus* StF 13038 could thus be attributed to excessive loss of larger droplets.

In emulsions, the lack of effective adsorption of cells onto the interface can be manifested by precipitation of cells and can often be accompanied by inadequate stability. Nevertheless, cell precipitation can also encounter in apparently stable emulsions, such as *Lentilactobacillus buchneri* DSM 20057, suggesting that it is multifactorial. For instance, some of the cells added to the emulsions, i.e., OD_600_ >2.0, could remain unbound or loosely bound to the interface, eventually undergoing precipitation over the course of storage, as was observed in many of the emulsions produced in this study. The inherent tendency of some cells to undergo rapid sedimentation can accelerate the rate of cell precipitation, apparently leading to formation of large cell precipitate, as was typically observed for emulsions of *Lb. crispatus* DSM 20584 and *Lb. hamsteri* CCUG 30722. In ideal circumstances, effective and irreversible adsorption of cells to the interface will have to diminish loss of cells due to precipitation. Observation of minimal cell precipitation concomitantly with remarkable stability of relative emulsion volumes for *Lb. acidophilus* StF 13015 and *Levilactobacillus brevis* CCUG 30670 is thus suggestive of irreversible interface adsorption of these two strains and the resultant stable Pickering emulsions.

In previous studies, the presence of unbound or loosely bound cells in the emulsions were revealed by detection of minor peaks on the DSD plots (Jiang et al., 2019). Virtually all of the distribution plots generated from our emulsions encompassed minor peaks in the range of 1-15 µm. Importantly, this size range is consistent with the cell sizes of rod-shaped LAB strains reported previously [36]. The sample to sample variation in the volume density of the minor peaks can thus be reflective of corresponding variation in the extent of bacterial adsorption to the interface. Observation of mostly less pronounced minor peak can thus signify low concentration of unbound or loosely bound cells in the emulsified sample fraction, thereby indicating successful interface adsorption.

The findings from d(4,3) measurement suggest that 10 strains were capable of forming adequately stable Pickering macro-emulsions, thus corroborating the existence of intrinsic Pickering stabilization potential among strains of ex-*Lactobacillus* species. The d(4,3) values (Figure 3) mostly overlapped with previously reported Pickering emulsions of ex-*Lactobacillus* species and strains [15], [17]. Nonetheless, not all the strains were equally outstanding with respect to formation of sufficiently small emulsion droplets. To this end, *Lb. acidophilus* StF 13015 appeared to be the most promising, forming the smallest droplets [d(4,3) of 61.42±25.67 µm to 67.73±3.21 µm], as compared to the rest of the group [d(4,3) of 84.62±6.70 to 88.32±5.29 µm]. The low precipitation of cells, coupled with detection of negligible minor peak on the distribution plot, substantiates that this strain could establish effective interface adsorption. Given the potential of this strain to enhance emulsion volume, generate fine gross texture and extremely firm consistency, and promote adequate physical and droplet stability, it is appealing to conclude that this strain is among the most promising bacterial Pickering stabilizers.

## 5. Conclusion

A collection of naturally occurring ex-*Lactobacillus* strains was characterized in relation to cell hydrophobicity and ZP measurement. Cell hydrophobicity was observed to be a natural phenomenon for a significant fraction of the tested strains. This contrasts with previous notions suggesting that most ex-*Lactobacillus* strains are intrinsically hydrophilic. A significant fraction of these strains appears to have intrinsic propensity to serve as Pickering stabilizers of emulsions and possibly other colloidal food materials. Such hydrophobic cells were observed to generate stable emulsions. This was confirmed using various droplet size measurement techniques.

---

**Figure S1.**
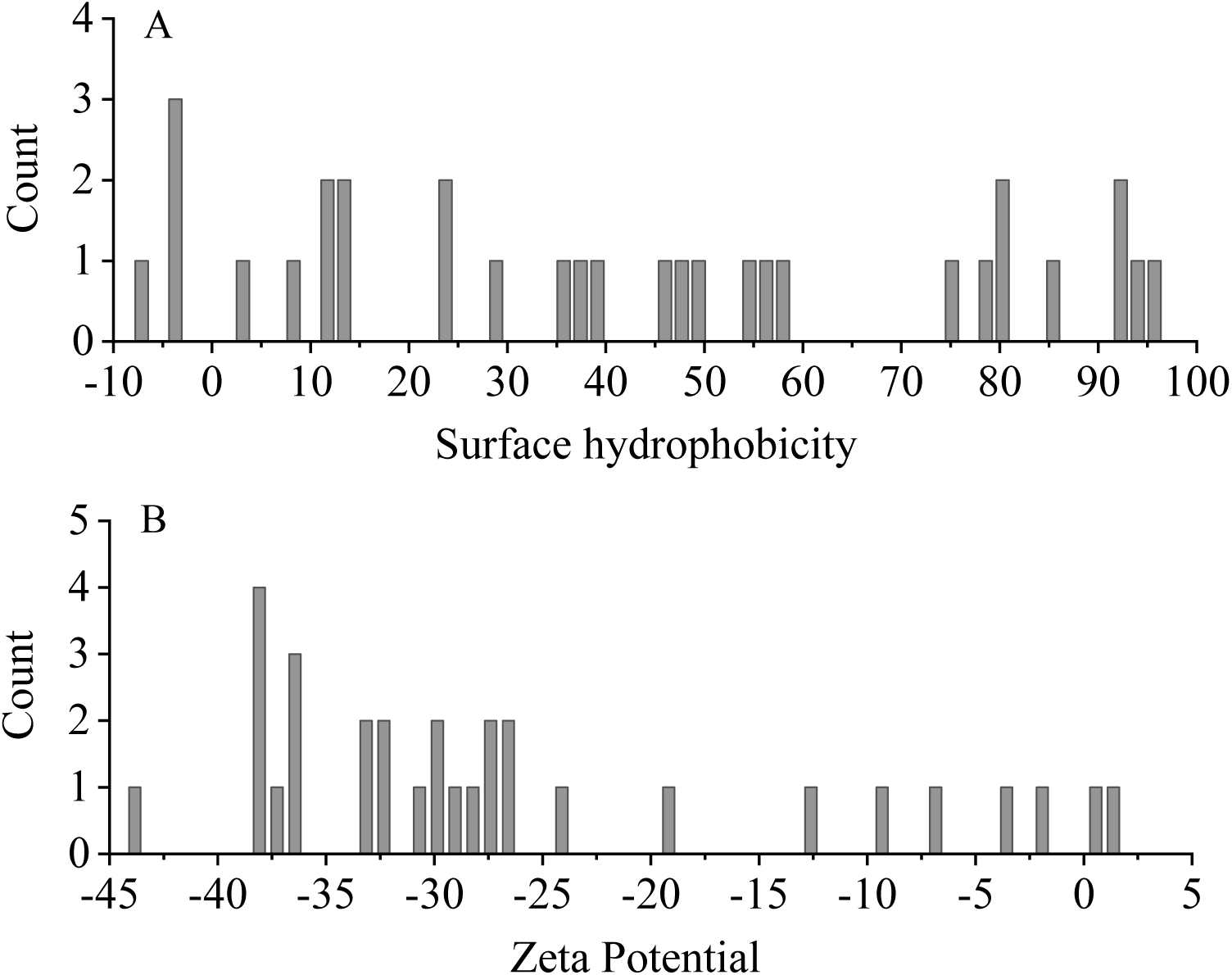
Distribution of ex-*Lactobacillus* strains at certain value intervals of (B) zeta potential and (A) surface hydrophobicity, corresponding to bin sizes of 0.82 and 1.71, respectively.

**Figure S2.**
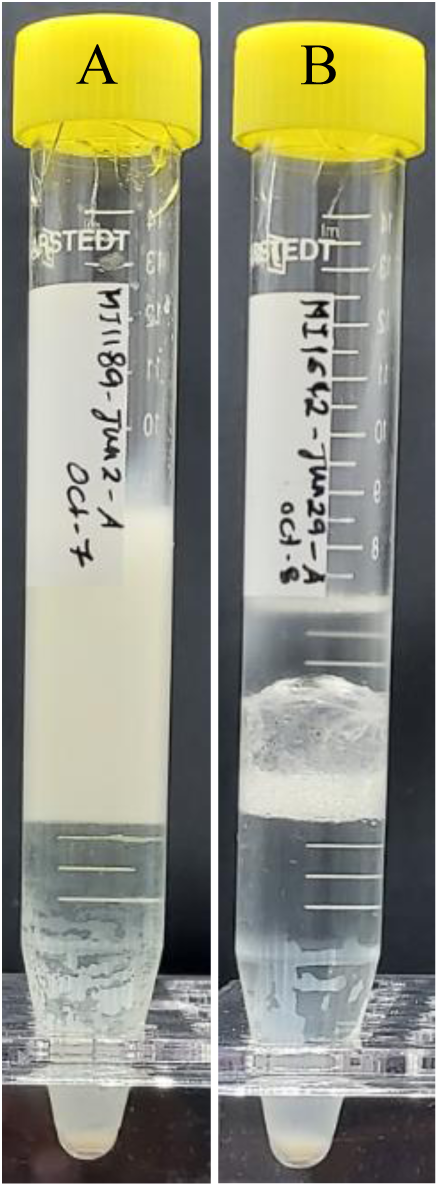
Pickering emulsions induced by (A) *Lb. acidophilus* StF 13015 and (B) *Lentilactobacillus parabuchneri* CCUG 32261 after prolonged storage (2 months), showing relatively stable and less stable emulsions, respectively, according to macroscopic appearances.

**Figure S3.**
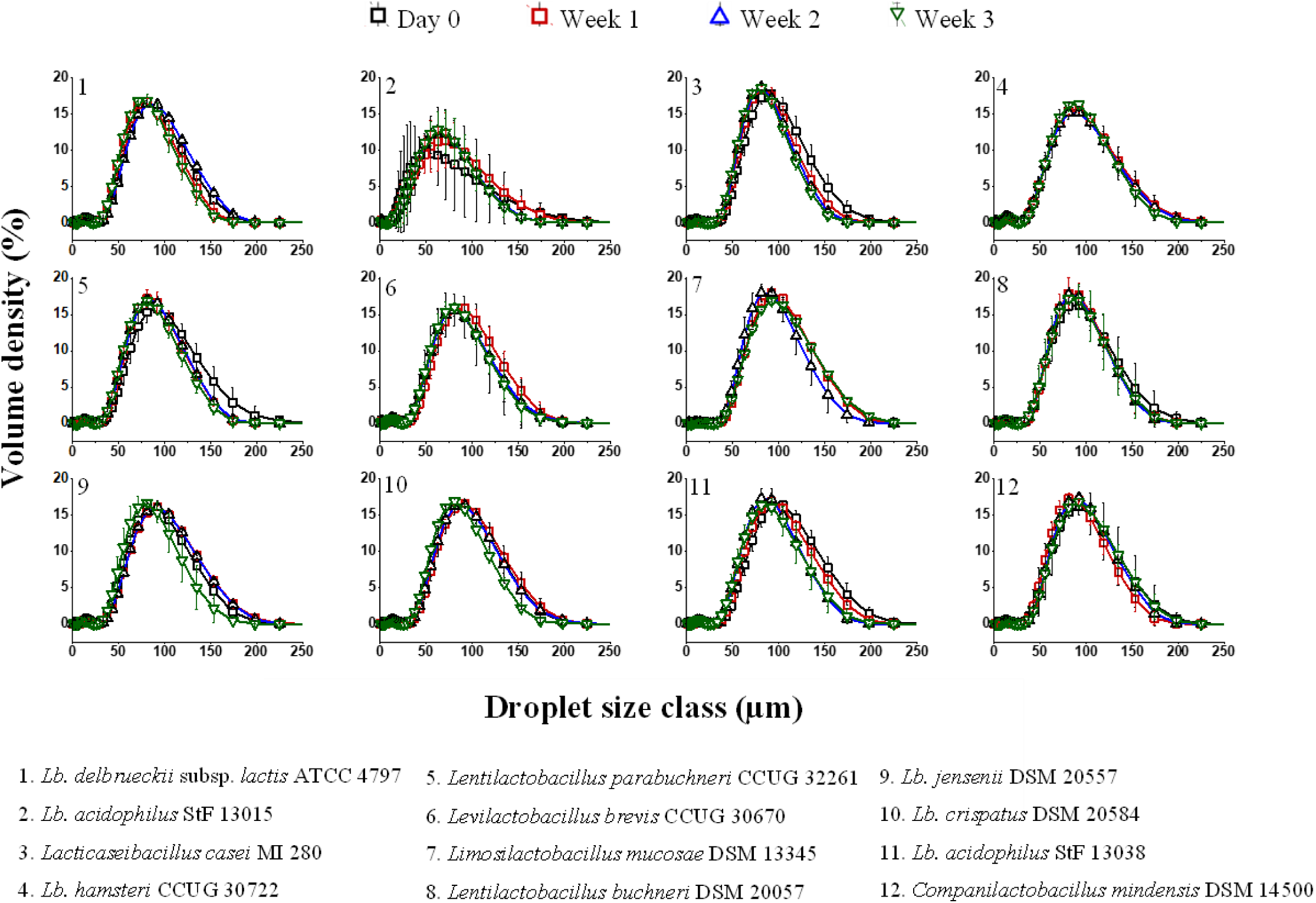
Droplet size distributions of Pickering emulsions on the day of preparation (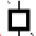), and after storage for 1 (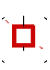), 2 (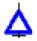) and 3 (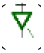) weeks.

**Figure S4.**
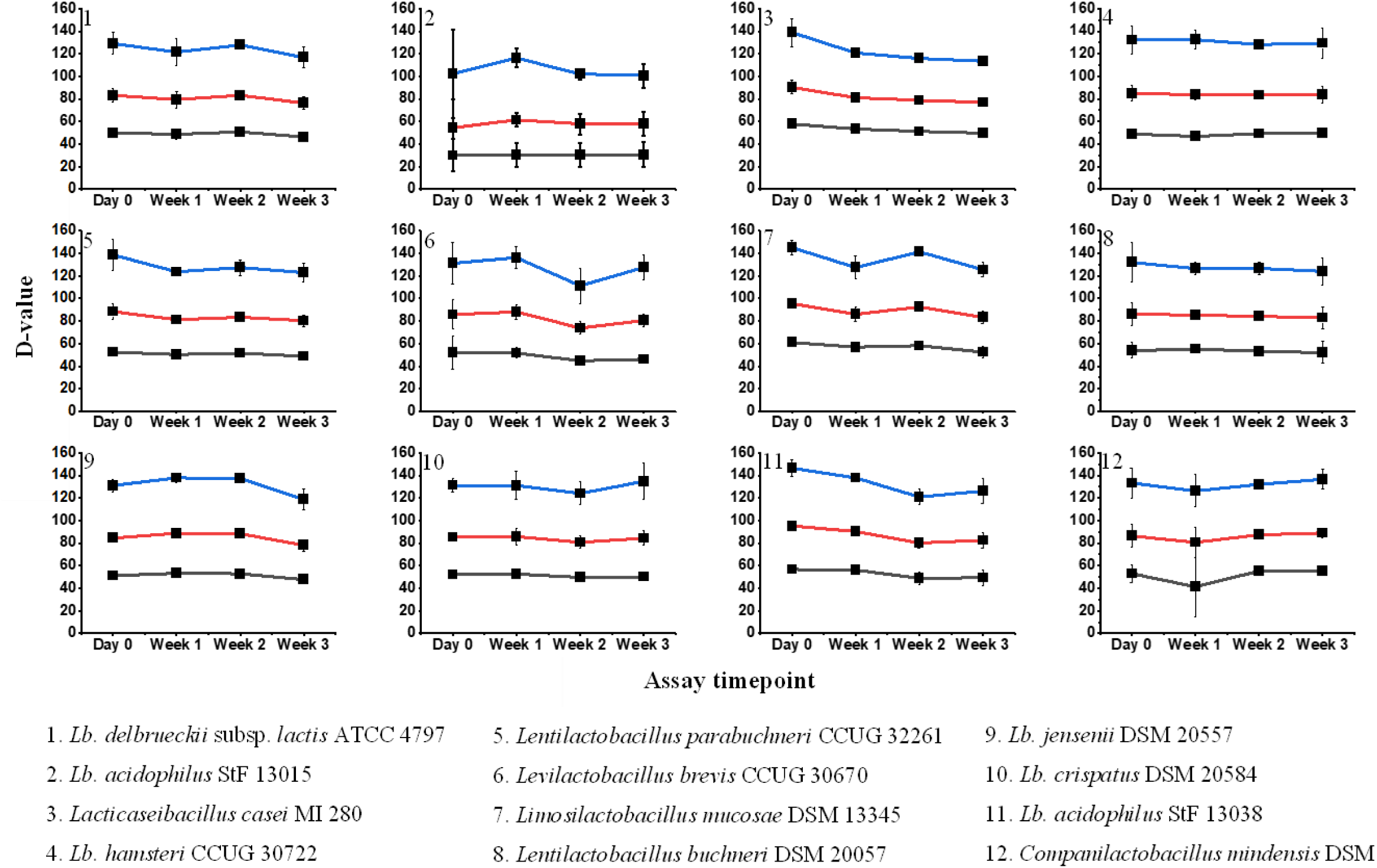
Droplet diameters of emulsions at 10% [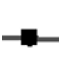], 50% [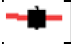] and 90% [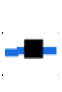] cumulative volumes, measured on the day of preparation and at various storage times.

**Figure S5.**
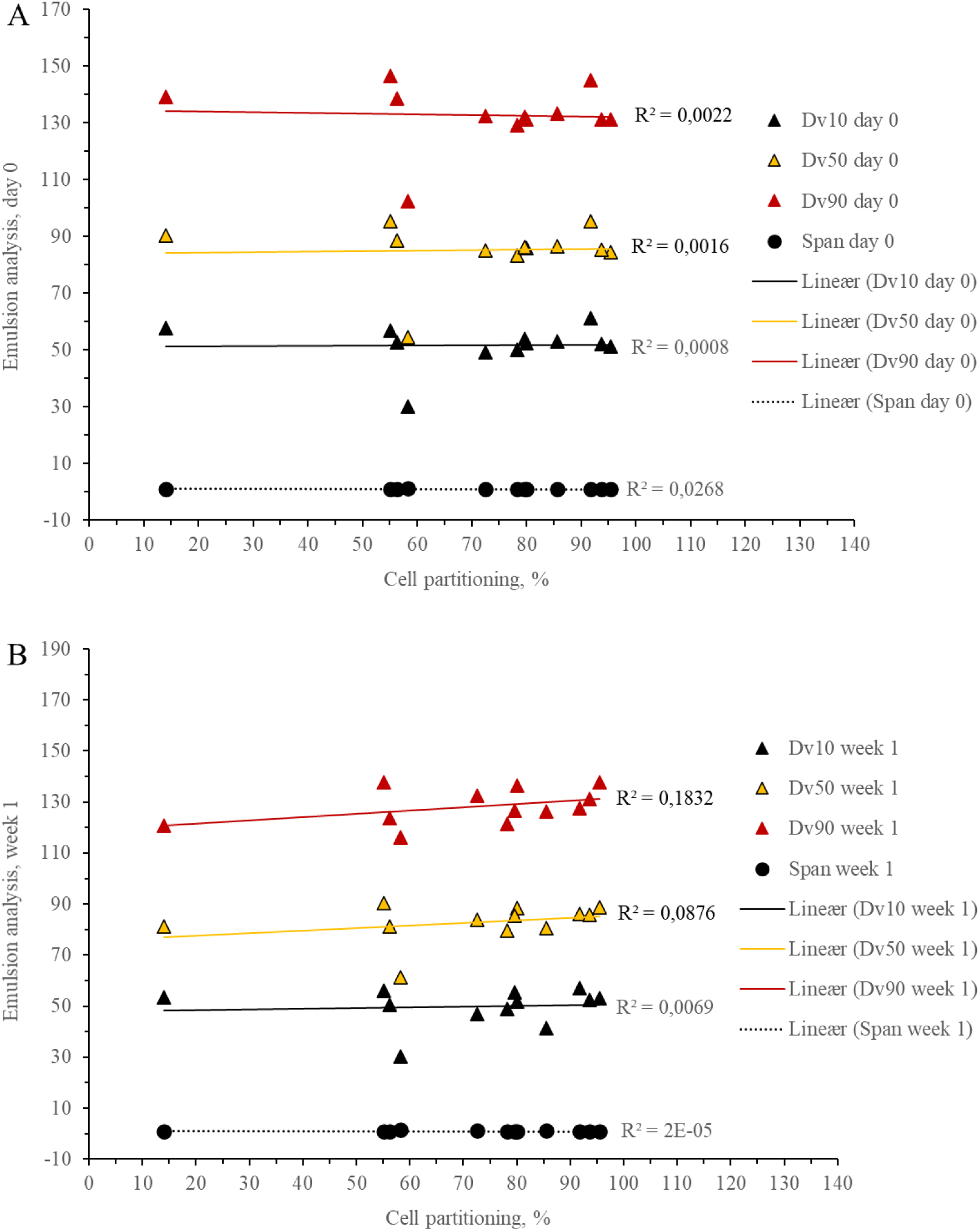
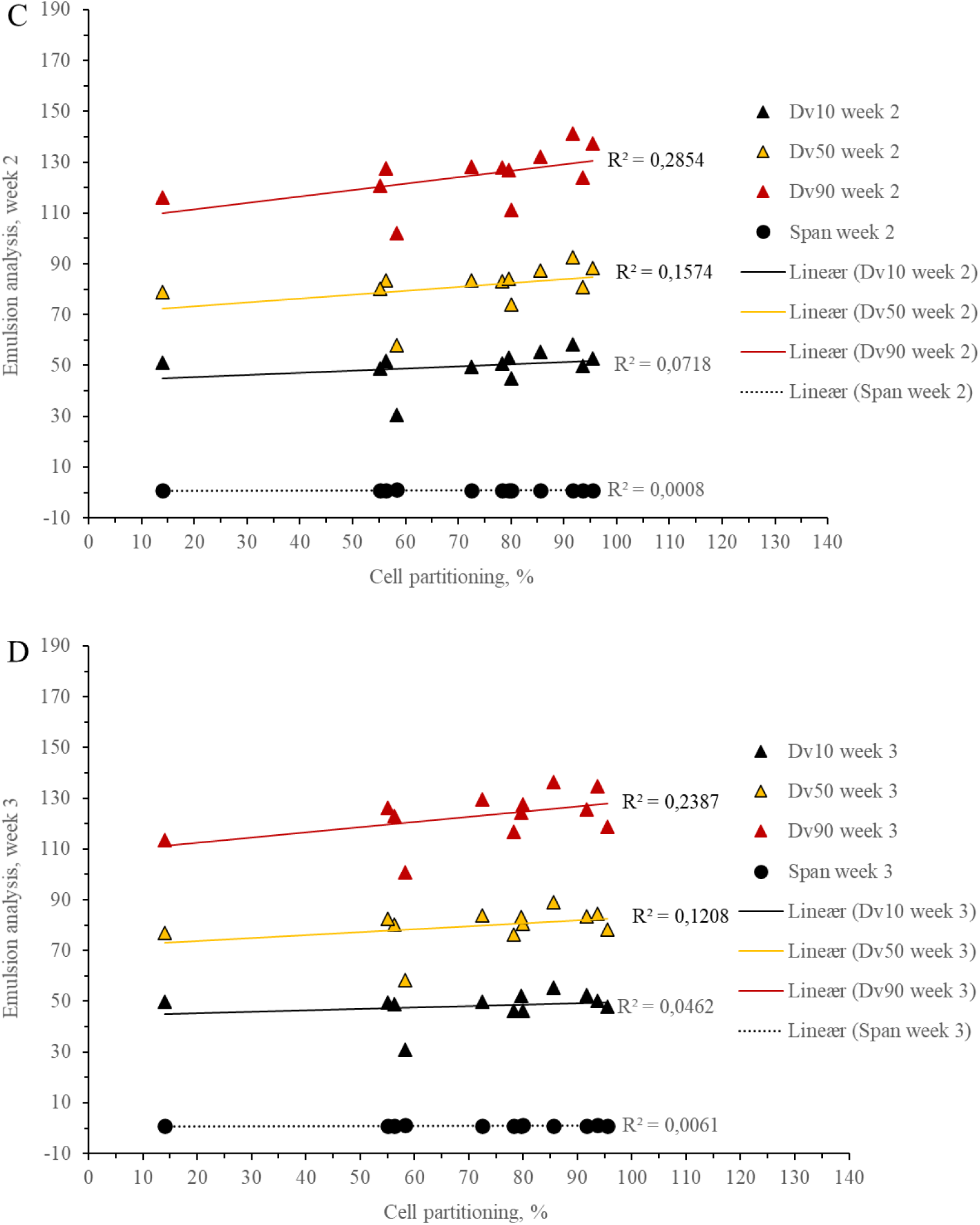
Correlation plots for MATH vs Dv10, Dv50, Dv90 (droplet diameters at 10%, 50% and 90% cumulative volumes, respectively) and relative span for day 0 (A), week 1 (B), 2 (C) and 3 (D). Cell partitioning does not show linear correlation with D values and span.

**Figure S6.**
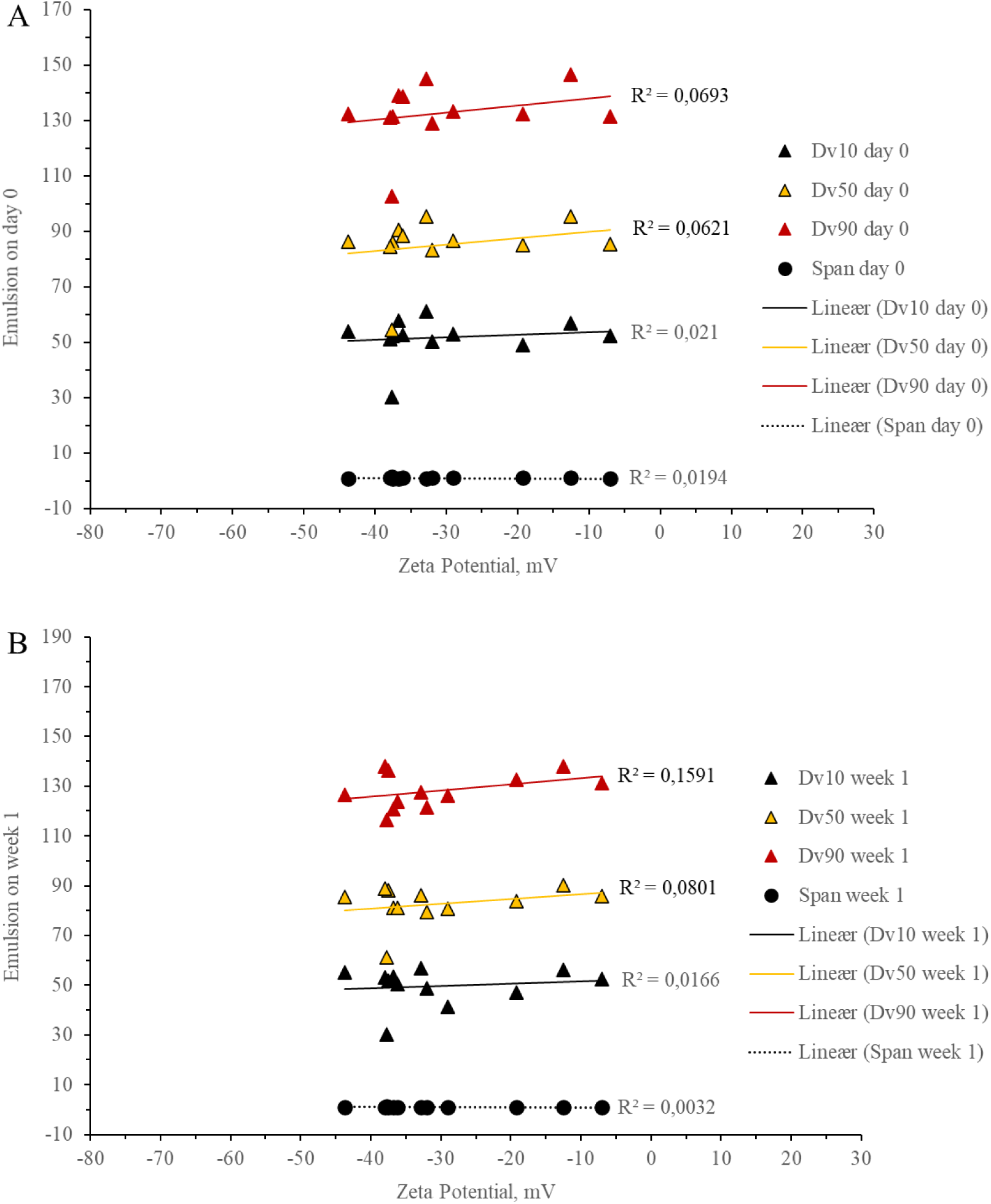
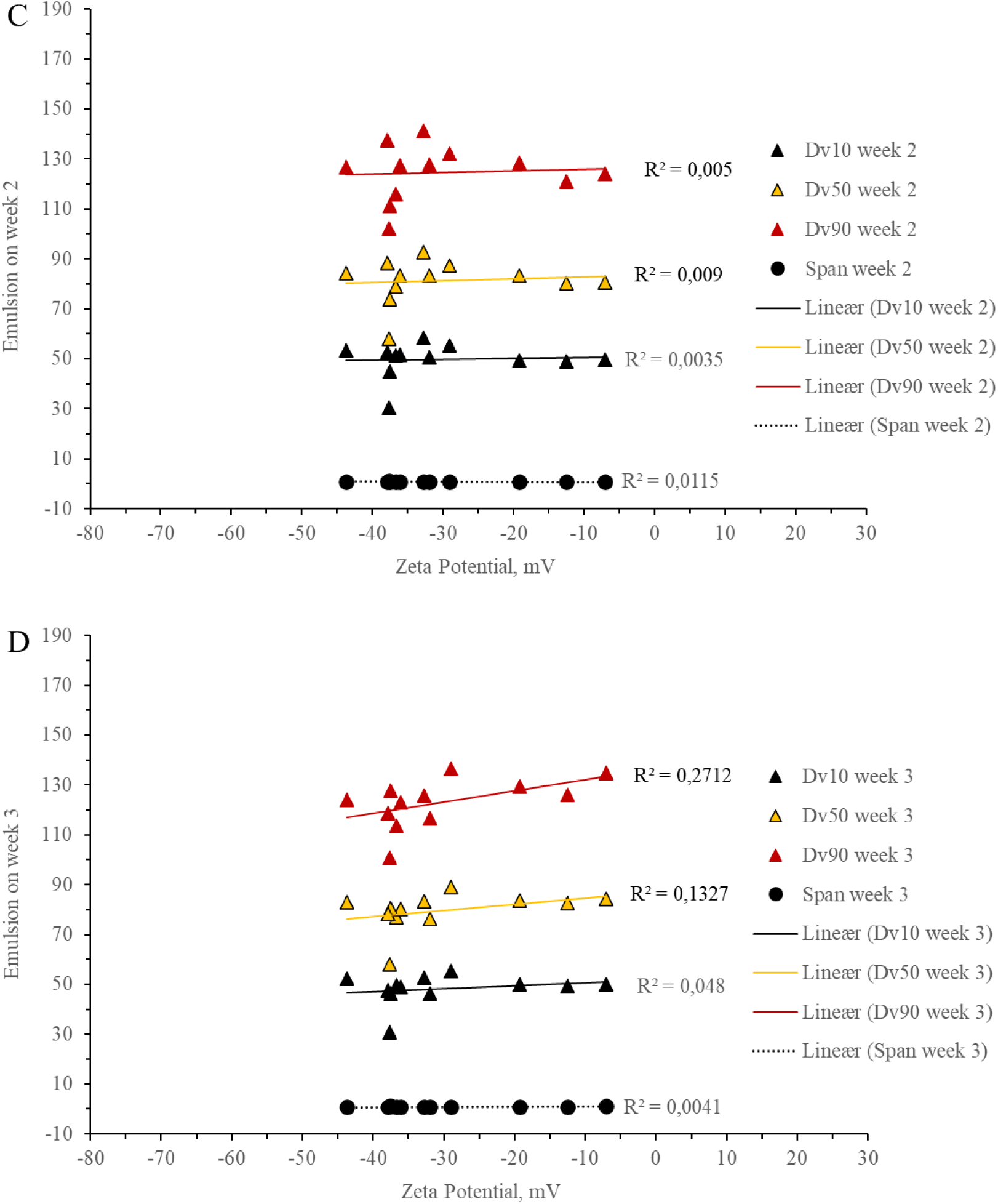
Correlation plots for ZP vs Dv10, Dv50, Dv90 (droplet diameters at 10%, 50% and 90% cumulative volumes, respectively) and relative span for day 0 (A), week 1 (B), 2 (C) and 3 (D). Cell partitioning does not show linear correlation with D values and span.

**Figure S7.**
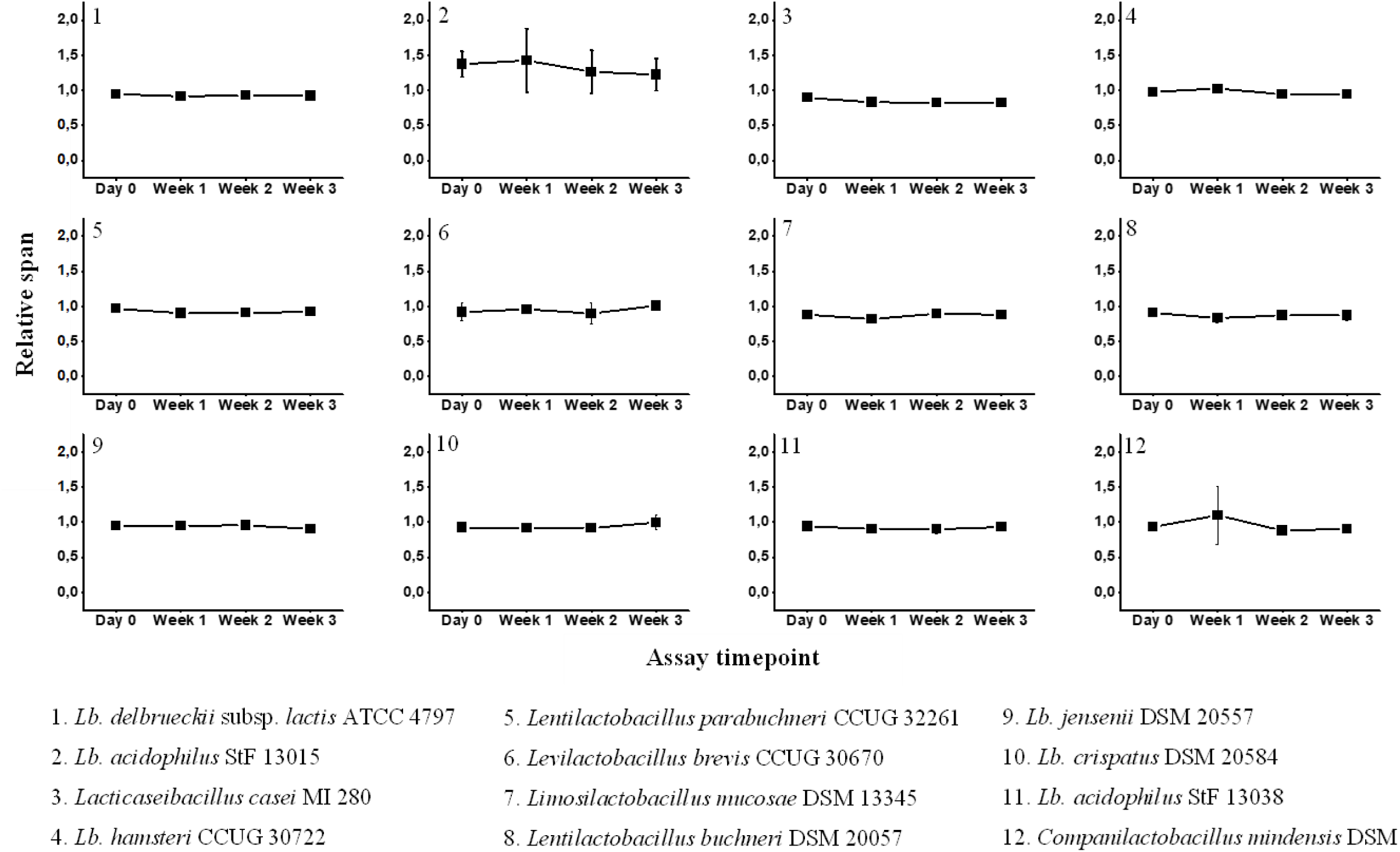
The width of the distribution of the droplet sizes of oil-in-water emulsions at various storage

**Table S1.**
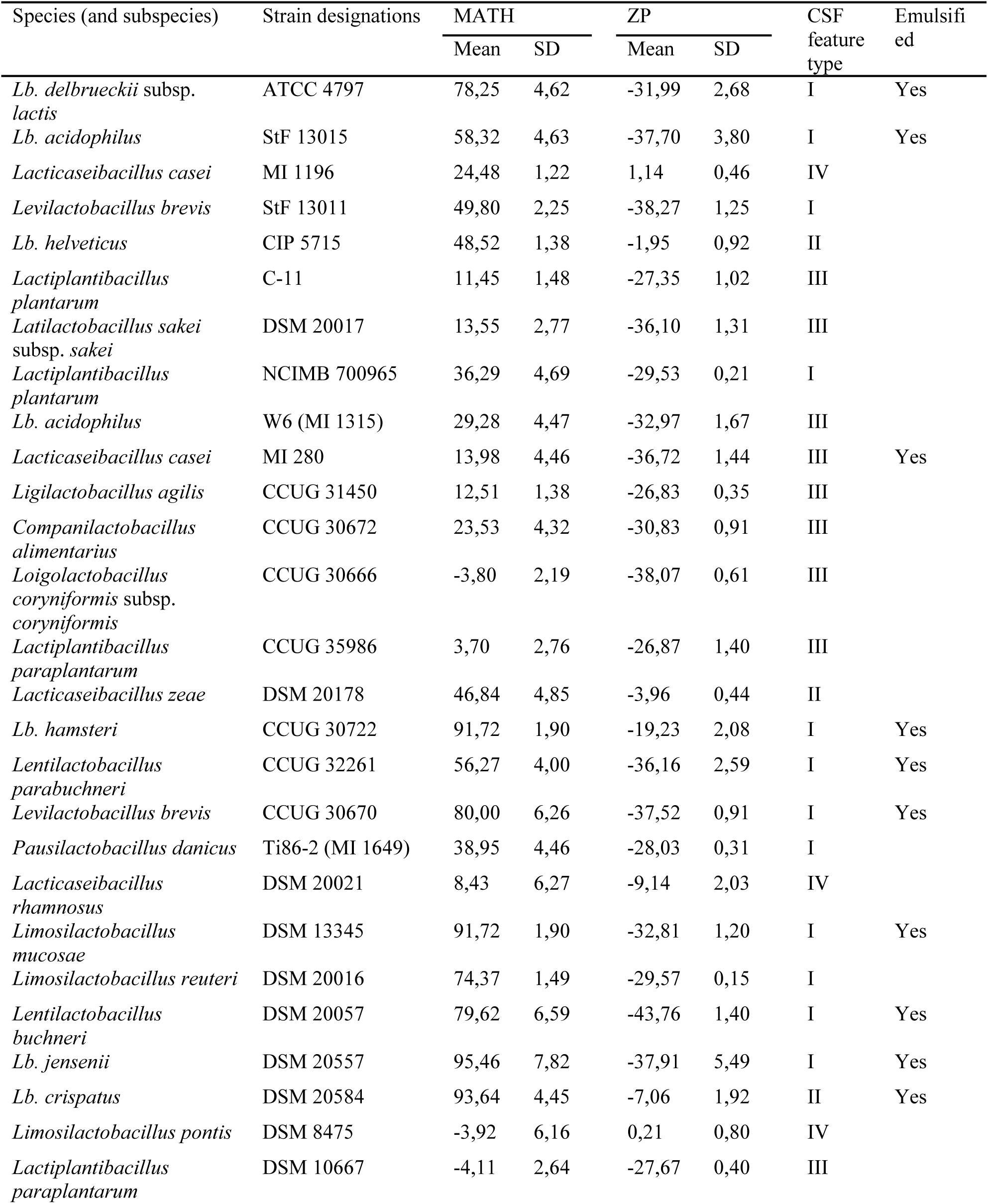
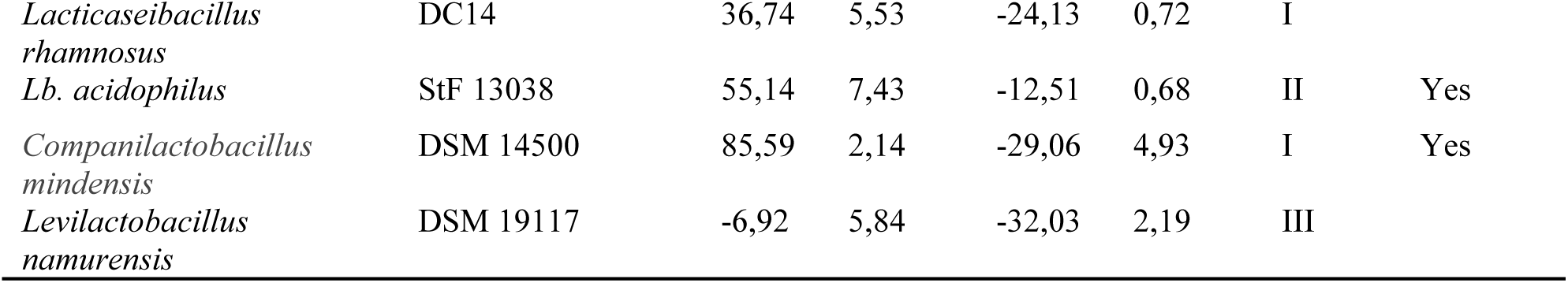
MATH and ZP values, and cell surface features of ex-*Lactobacillus* strains

| Species (and subspecies) | Strain designations | MATH |  | ZP |  | CSF<br>feature<br>type | Emulsifi<br>ed |
| --- | --- | --- | --- | --- | --- | --- | --- |
|  |  | Mean | SD | Mean | SD |  |  |
| *Lb. delbrueckii* subsp. *lactis* | ATCC 4797 | 78,25 | 4,62 | -31,99 | 2,68 | I | Yes |
| *Lb. acidophilus* | StF 13015 | 58,32 | 4,63 | -37,70 | 3,80 | I | Yes |
| *Lacticaseibacillus casei* | MI 1196 | 24,48 | 1,22 | 1,14 | 0,46 | IV |  |
| *Levilactobacillus brevis* | StF 13011 | 49,80 | 2,25 | -38,27 | 1,25 | I |  |
| *Lb. helveticus* | CIP 5715 | 48,52 | 1,38 | -1,95 | 0,92 | II |  |
| *Lactiplantibacillus plantarum* | C-11 | 11,45 | 1,48 | -27,35 | 1,02 | III |  |
| *Latilactobacillus sakei* subsp. *sakei* | DSM 20017 | 13,55 | 2,77 | -36,10 | 1,31 | III |  |
| *Lactiplantibacillus plantarum* | NCIMB 700965 | 36,29 | 4,69 | -29,53 | 0,21 | I |  |
| *Lb. acidophilus* | W6 (MI 1315) | 29,28 | 4,47 | -32,97 | 1,67 | III |  |
| *Lacticaseibacillus casei* | MI 280 | 13,98 | 4,46 | -36,72 | 1,44 | III | Yes |
| *Ligilactobacillus agilis* | CCUG 31450 | 12,51 | 1,38 | -26,83 | 0,35 | III |  |
| *Companilactobacillus alimentarius* | CCUG 30672 | 23,53 | 4,32 | -30,83 | 0,91 | III |  |
| *Loigolactobacillus coryniformis* subsp. *coryniformis* | CCUG 30666 | -3,80 | 2,19 | -38,07 | 0,61 | III |  |
| *Lactiplantibacillus paraplantarum* | CCUG 35986 | 3,70 | 2,76 | -26,87 | 1,40 | III |  |
| *Lacticaseibacillus zeae* | DSM 20178 | 46,84 | 4,85 | -3,96 | 0,44 | II |  |
| *Lb. hamsteri* | CCUG 30722 | 91,72 | 1,90 | -19,23 | 2,08 | I | Yes |
| *Lentilactobacillus parabuchneri* | CCUG 32261 | 56,27 | 4,00 | -36,16 | 2,59 | I | Yes |
| *Levilactobacillus brevis* | CCUG 30670 | 80,00 | 6,26 | -37,52 | 0,91 | I | Yes |
| *Pausilactobacillus danicus* | Ti86-2 (MI 1649) | 38,95 | 4,46 | -28,03 | 0,31 | I |  |
| *Lacticaseibacillus rhamnosus* | DSM 20021 | 8,43 | 6,27 | -9,14 | 2,03 | IV |  |
| *Limosilactobacillus mucosae* | DSM 13345 | 91,72 | 1,90 | -32,81 | 1,20 | I | Yes |
| *Limosilactobacillus reuteri* | DSM 20016 | 74,37 | 1,49 | -29,57 | 0,15 | I |  |
| *Lentilactobacillus buchneri* | DSM 20057 | 79,62 | 6,59 | -43,76 | 1,40 | I | Yes |
| *Lb. jensenii* | DSM 20557 | 95,46 | 7,82 | -37,91 | 5,49 | I | Yes |
| *Lb. crispatus* | DSM 20584 | 93,64 | 4,45 | -7,06 | 1,92 | II | Yes |
| *Limosilactobacillus pontis* | DSM 8475 | -3,92 | 6,16 | 0,21 | 0,80 | IV |  |
| *Lactiplantibacillus paraplantarum* | DSM 10667 | -4,11 | 2,64 | -27,67 | 0,40 | III |  |
| *Lacticaseibacillus*<br>*rhamnosus* | DC14 | 36,74 | 5,53 | -24,13 | 0,72 | I |  |
| *Lb. acidophilus* | StF 13038 | 55,14 | 7,43 | -12,51 | 0,68 | II | Yes |
| *Companilactobacillus*<br>*mindensis* | DSM 14500 | 85,59 | 2,14 | -29,06 | 4,93 | I | Yes |
| *Levilactobacillus*<br>*namurensis* | DSM 19117 | -6,92 | 5,84 | -32,03 | 2,19 | III |  |

**Table S2.** Relative emulsion volumes one week after storage of the samples at room temperature

| Selected strain | ERV, % |  |
| --- | --- | --- |
|  | Mean | sd |
| *Lb. acidophilus* StF 13015 | 67,57 | 3,82 |
| *Limosilactobacillus mucosae* DSM 13345 | 66,57 | 2,23 |
| *Lentilactobacillus parabuchneri* CCUG 32261 | 59,38 | 13,26 |
| *Lentilactobacillus buchneri* DSM 20057 | 58,76 | 11,19 |
| *Lacticaseibacillus casei* MI 280 | 58,33 | 11,79 |
| *Lb. crispatus* DSM 20584 | 57,14 | 0,00 |
| *Lb. acidophilus* StF 13038 | 56,40 | 9,47 |
| *Lb. jensenii* DSM 20557 | 56,28 | 1,50 |
| *Lb. hamsteri* CCUG 30722 | 50,00 | 3,33 |
| *Lb. delbrueckii* subsp. *lactis* ATCC 4797 | 46,62 | 4,78 |
| *Companilactobacillus mindensis* DSM 14500 | 44,13 | 3,89 |
| *Levilactobacillus brevis* CCUG 30670 | 41,67 | 11,79 |

## Notes

### Competing Interest Statement

The authors have declared no competing interest.

